# The adaptive response to long-term nitrogen starvation in *Escherichia coli* requires the breakdown of allantoin

**DOI:** 10.1101/2020.03.30.016519

**Authors:** Amy Switzer, Lynn Burchell, Josh McQuail, Sivaramesh Wigneshweraraj

## Abstract

Bacteria initially respond to nutrient starvation by eliciting large-scale transcriptional changes. The accompanying changes in gene expression and metabolism allow the bacterial cells to effectively adapt to the nutrient starved state. How the transcriptome subsequently changes as nutrient starvation ensues is not well understood. We used nitrogen (N) starvation as a model nutrient starvation condition to study the transcriptional changes in *Escherichia coli* experiencing long-term N starvation. The results reveal that the transcriptome of N starved *E. coli* undergoes changes that are required to maximise chances of viability and to effectively recover growth when N starvation conditions become alleviated. We further reveal that, over time, N starved *E. coli* cells rely on the degradation of allantoin for optimal growth recovery when N becomes replenished. This study provides insights into the temporally coordinated adaptive responses that occur in *E. coli* experiencing sustained N starvation.

**IMPORTANCE:** Bacteria in their natural environments seldom encounter conditions that support continuous growth. Hence, many bacteria spend the majority of their time in states of little or no growth due to starvation of essential nutrients. To cope with prolonged periods of nutrient starvation, bacteria have evolved several strategies, primarily manifesting themselves through changes in how the information in their genes is accessed. How these coping strategies change over time under nutrient starvation is not well understood and this knowledge is not only important to broaden our understanding of bacterial cell function, but also to potentially find ways to manage harmful bacteria. This study provides insights into how nitrogen starved *Escherichia coli* bacteria rely on different genes during long term nitrogen starvation.

## INTRODUCTION

The process of RNA synthesis, transcription, is often the first decision-making and regulatory point in the flow of genetic information and is catalysed by the multisubunit enzyme, RNA polymerase (RNAP) (1). The transcriptional responses that establish and maintain specific cellular states are dynamic and are often controlled by many transcription regulatory factors that influence the activity of the RNAP (1). Conditions that sustain constant bacterial growth are seldom found in nature. Whether in a host or the environment, bacteria frequently encounter nutrient deprivation, which can attenuate bacterial growth (2-4). Hence, the growth attenuated state is now widely acknowledged as an important physiological state in bacterial cell function. However, although the transcriptional basis of bacterial growth is widely studied, our understanding of the transcriptome of growth attenuated bacteria remains in its infancy. In fact, several recent studies on different species of bacteria have revealed that growth attenuated bacteria are transcriptionally active. For example: Gefen et al showed that a batch culture of *E. coli* cells retain transcriptional activity in the stationary phase of growth over a period of nearly 2 days (5); Stapels et al showed that non-growing intracellular *Salmonella* in host macrophages are transcriptionally active and this activity is important to shape the host environment to benefit bacterial survival (6); similarly, non-replicating *Mycobacterium tuberculosis* cells retained transcriptional activity in the lungs of mice (7); and more recently, Gray et al reported that *Bacillus subtilis* cells under oligotrophic conditions are transcriptionally active and replicate at an extremely slow rate (8). Therefore, understanding the transcriptional landscape in growth attenuated bacteria is important as this knowledge could inform and inspire new antibacterial strategies, such as disrupting genetic networks and products that are activated during growth attenuation, which subsequently become essential for survival and/or optimal growth recovery.

In this study, we used nitrogen (N) starvation as a model condition to understand how the transcriptional landscape in nutrient starvation-induced growth attenuated *E. coli* affects bacterial survival and ability to recover growth when nutrients (i.e. N) become replenished. N is an integral nutrient to bacterial cell function since N is an essential element of most macromolecules in the bacterial cell (proteins, metabolites, nucleic acids and cell wall components). Therefore, an exponentially growing batch culture of *E. coli* rapidly attenuates growth upon N run-out (9-11). The initial adaptive response to N run-out, the N regulation (Ntr) stress response, results in the large scale and multilevel reprogramming of the transcriptome, involving ∼40% of all *E. coli* genes, to allow *E. coli* to adjust to N starvation (12-14). Since *E. coli* perceive N run-out as a decrease in the internal concentration of glutamine, the genes expressed as part of the Ntr response are predominantly involved in the transport and assimilation of nitrogenous compounds into glutamine or glutamate, yielding these catabolically or sparing the requirement for them in cellular processes (12, 13). Although our understanding of the initial adaptive transcriptional response to N starvation is well established, how the transcriptome changes during long-term N starvation is far from clear.

## RESULTS AND DISCUSSION

### Inhibition of transcription in N starved *E. coli* compromises survival and growth recovery

To setup an experimental system to study how the transcriptome changes during long-term N starvation, we grew a batch culture of *E. coli* in a defined minimal growth medium in the presence of a limiting amount (3 mM) of ammonium chloride as the sole N source and excess (22 mM) glucose as the carbon (C) source. As shown in Fig. 1A (left and middle panels), under these conditions, when ammonium chloride becomes depleted (but sufficient glucose is still available to support growth), bacteria attenuate growth due to N starvation (12). From the onset of N starvation (referred to as N-), the number of cells in the population of growth attenuated *E. coli*, as measured by the number of colony forming units (c.f.u.) in the population, increases at a slow rate over the initial 24 h under N starvation (doubling time ∼39 h, which is ∼80-fold less than during exponential growth when N is abundant; referred to as N+) (Fig. 1A, right panel, orange area). This growth seems to occur despite the absence of any detectable ammonium chloride in the growth medium (Fig. 1A, left and middle panels). However, as N starvation ensues beyond 24 h (referred to as N-24), the population ceases growth before the number of viable cells gradually begins to decline after 48 h under N starvation (Fig. 1A, red area). Therefore, the first 24 h period following N-, which we refer to as the N starved adaptive phase (NSAP), seems to be an important adaptive period during long-term N starvation. It is likely that transcriptional changes occur during NSAP and we sought to determine how the inhibition of transcription during NSAP affects cell survival and growth recovery when N starvation stress is alleviated. To do this, we treated the bacteria with 150 μg/ml of the bacteriostatic antibiotic rifamycin (Rif) at ∼1 h following N-for ∼23 h. As previously shown by us, under these conditions, 150 μg/ml Rif prevents transcription elongation by the RNAP by trapping it at promoters (12). We initially compared the proportion of viable cells in the Rif treated and untreated population of bacteria at N-24 (i.e. ∼23 h following Rif treatment at N-1). As shown in Fig. 1B, at N-24, only ∼50% cells in the Rif treated culture were viable compared to the Rif untreated culture. Rif treatment also had a profound effect on the ability of bacteria from N-24 to resume growth compared to untreated bacteria. As shown in Fig. 1C and 1D, the lag time to growth recovery of the Rif treated bacteria (following c.f.u. correction to ensure an equal number of viable treated and untreated bacteria were used; based on data in Fig. 1B) from N-24 was delayed by ∼4.8 h compared to untreated bacteria. We considered whether this difference in lag time between Rif treated and untreated bacteria was due to the presence of a slower growing subpopulation(s) of bacteria in the Rif treated population that resumed growth slowly. To investigate this, we used a method called ScanLag (15), which allows measurement of appearance time of individual colonies on N replete solid growth media as a function of time following inoculation. We expected that if the Rif treated population consisted of one or more subpopulations compared to the untreated population, then the appearance of individual colonies of bacteria treated with Rif would be more heterogenous than that of untreated bacteria. As shown in Fig. 1E, no heterogeneity in growth in the populations of Rif treated and untreated bacteria from N-24 was detected. However, the ScanLag data substantiated the data in Fig. 1C and 1D and demonstrated that Rif treated cells have an increased lag time to growth recovery compared to untreated cells. Once growth resumed, the doubling time of Rif treated, and untreated bacteria did not markedly differ (Fig. 1D). Overall, we conclude that inhibition of transcription at the onset of NSAP has a detrimental effect on the ability of *E. coli* to adapt to long-term N starvation in terms of maximising chances of viability and optimal growth recovery when N becomes available.

**Fig 1.**
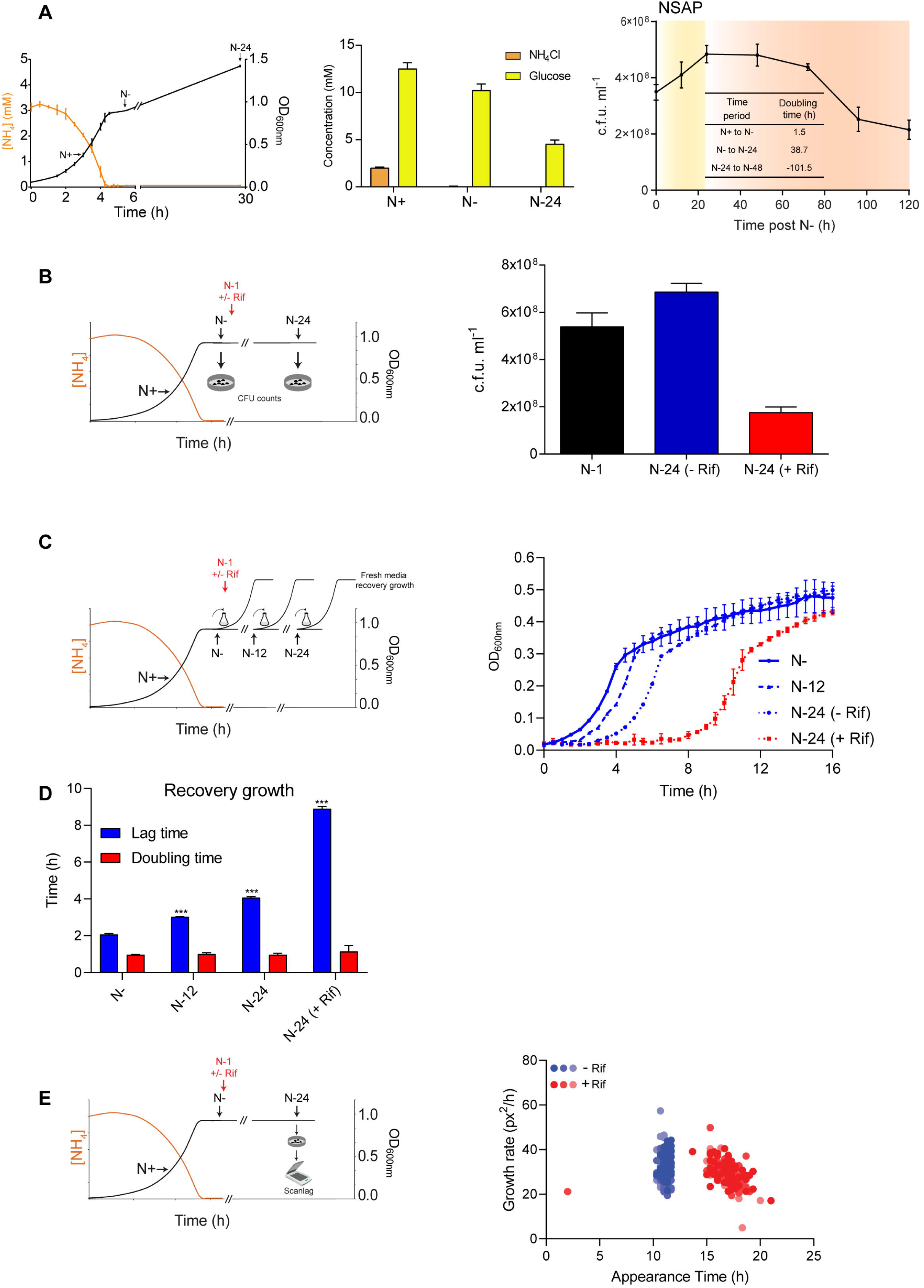
Inhibition of transcription in N starved *E. coli* compromises survival and growth recovery. (A) Graph showing a representative growth curve (measured by OD_600nm_ readings) of *E. coli* and concentration (mM) of ammonium chloride in the growth medium as a function of growth (left panel). The time points referred to as N+, N- and N-24 in the text are indicated on the growth curve. The bar graph shows the concentration (mM) of ammonium chloride and glucose at N+, N- and N-24 (middle panel). Graph showing viability, measured by counting colony forming units (c.f.u.) of *E. coli* over 5 days under N starvation (right panel). Inset table shows doubling time between different growth periods and the NSAP is indicated. (B) Bar graph showing the proportion of viable cells (measured by c.f.u.) in the population of *E. coli* at N-1 (black), following 24 h under N starvation with (red) and without (blue) Rif treatment at N-1. (C) Graph showing recovery growth in fresh N replete medium as measured by OD_600nm_ readings. The time points and conditions from which bacteria were sampled for recovery growth experiments are indicated and discussed in the text. (D) Graph showing lag time and doubling time during growth recovery experiments from (C). (E) ScanLag analysis of colony growth rate and appearance time after plating of 24 h N starved bacteria on LB agar after Rif treatment (red) and no Rif treatment (blue) at N-1. The various shades of the same colour represent distinct biological replicates, where n=3. In (A)-(D), error bars represent SEM, where n=3 and the schematics describe the different experiments for ease of understanding.

### The transcriptome *E. coli* undergoes changes during the NSAP

To study the transcriptional changes that occur during the NSAP, we compared, as indicated in Fig. 2A, the transcriptomes of *E. coli* during exponential growth (N+), following initial N run-out (N-), and at N-24. We defined differentially expressed genes as those with expression levels changed ≥ 2-fold with a false discovery rate adjusted *P*-value < 0.05. As previously reported by us and others, the initial transcriptional response to N starvation involves differential expression of 1787 genes primarily required for scavenging and catabolism of alternative N sources (13, 14, 16) and thus, we did not investigate these genes further (Phase A in Fig. 2A).

**Fig 2.**
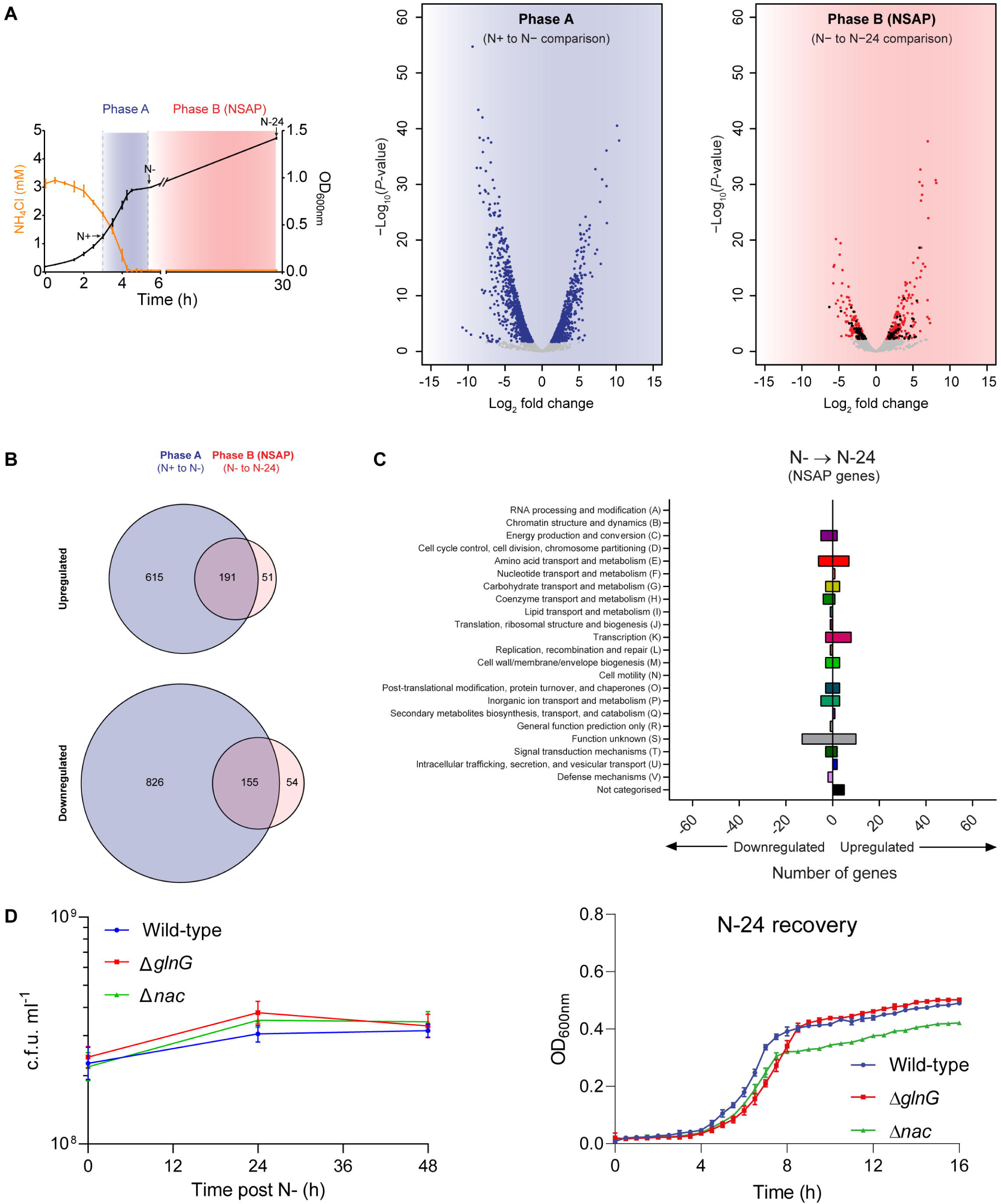
The transcriptome *E. coli* undergoes changes during the NSAP. (A) Graph showing a representative growth curve (measured by OD_600nm_ readings) of *E. coli* and concentration (mM) of ammonium chloride in the growth medium as a function of growth (left panel). The points at which samples were taken for RNA-seq analysis (N+, N- and N-24) and the phases referred to as A and B in the text are indicated. Graphs showing the log_2_ fold change in gene expression at N- compared to N+ (Phase A), and N-24 compared to N- (Phase B). Genes in blue (Phase A) or red (Phase B = NSAP) represent significant differential expression above a log_2_ fold change of 1 and a false discovery rate adjusted *P* value < 0.05 (right panel). Genes in black (Phase B) represent the NSAP genes, defined as having significant differential gene expression unique to Phase B (NSAP), but not Phase A. (B) Venn diagrams showing genes significantly upregulated (top) and downregulated (bottom) in Phase A compared to Phase B. (C) Graph showing Clusters of Orthologous Groups (COG) categorisation of NSAP genes, and the number of genes within each category whose expression is up- or downregulated from N-. (D) Graph showing viability as measured by c.f.u. counting of viable cells in the population of WT (blue), Δ*glnG* (red) and Δ*nac* (green) bacteria over 48 h of N starvation (left panel). Graph showing growth recovery of N-24 WT (blue), Δ*glnG* (red) and Δ*nac* (green) bacteria in N replete fresh media as measured by OD_600nm_ readings (right panel). In both graphs, error bars represent SEM, where n=3.

However, during the NSAP, upon transition from N- to N-24 (Phase B in Fig. 2A), 451 genes (∼10% of *E. coli* genes) are differentially expressed, of which 105 genes (∼2% of *E. coli* genes) become differentially expressed in Phase B, that were not differentially expressed in Phase A (Fig. 2B). Functional categorisation of the genes uniquely differentially expressed in Phase B, hereafter called NSAP genes, revealed that they are involved in diverse processes including transcription, amino acid transport, metabolism and several genes of unknown function (Fig. 2C). Eleven (*abgR, bfd, ileY, leuO, ompN, sibE, tauA, ycaD, yedL, yhdV*, and *yihO*) of the upregulated and nine (*evgA, glnA, gltI, hemH, ivy, pfkB, ybjX, yieP*, and *yjeJ*) of the downregulated NSAP genes were known to be regulated by either NtrC (*glnA* and *gltI*) or Nac (all other genes listed above) (16, 17) - the master transcription regulators that activate the initial adaptive response to N starvation. However, the absence of either *glnG* (which encodes for NtrC) or *nac* did not detectably affect viability of bacteria following 24 h under N starvation or the ability of 24 h N starved bacteria to recover growth when inoculated into fresh N replete minimal growth media (Fig. 2D). Overall, we conclude that the transcriptome of *E. coli* undergoes changes during the NSAP, and this is consistent with the finding above that 150 μg/ml of Rif treatment during the NSAP has a detrimental effect on the viability of the *E. coli* population following 24 h under N starvation and its ability to recover growth when N becomes replenished (Fig. 1B-D). It therefore seems that the adaptive transcriptional changes that occur during the NSAP happen subsequently to the initial adaptive changes to N starvation and do not depend on NtrC (and Nac), which coordinates the initial adaptive transcriptional response to N run-out.

### Four upregulated NSAP genes are important for optimal growth recovery from long-term N starvation

We next investigated how the upregulated NSAP genes affected viability and growth recovery of long-term N starved *E. coli*. Of the 51 upregulated NSAP genes, 44 are available in the Keio single gene knockout collection (18). Thus, we determined how the absence of any of the upregulated NSAP genes affected growth recovery following 24 under N starvation. To do this, we grew the library of upregulated NSAP gene mutant bacteria in minimal growth media containing excess (10 mM) ammonium chloride as the sole N source and measured cell density after 16 h of growth. The growth of the library of upregulated NSAP gene mutant strains was done in a plate reader incubator which allowed continuous monitoring of growth of each mutant strain simultaneously to allow better side-by-side comparison. The results shown in Fig. 3A indicate that absence of any of the NSAP genes did not markedly affect growth under N replete conditions. However, we cannot exclude the possibility that the relative dynamics of growth of the individual mutant strains could have differed from that of the wild-type (WT) strain over the 16 h period of time. We note that bacteria in which the tryptophan biosynthesis genes are deleted, lack the γ subunit of formate dehydrogenase-N (*fdnI*), or ferric rhodotorulic acid outer membrane transporter (*fhuE*), are unable to grow in our minimal growth media, most likely due to auxotrophy (19). Next, we reinoculated bacteria that grew under N replete conditions into fresh minimal growth media containing limiting, i.e. 3 mM, ammonium chloride and determined how the absence of the upregulated NSAP genes affected viability at N-24, i.e. following 24 h under N starvation. The results in Fig. 3B show that the viabilities of the upregulated NSAP gene mutants were similar to that of the WT strain. When the 24 h N starved mutant bacteria were inoculated into fresh N replete minimal growth media, the dynamics of growth recovery for the majority of the mutant strains were similar to that of the WT strain (Fig. 3C). However, Δ*allB*, Δ*leuO*, Δ*ynfP*, and Δ*ybcJ* mutant strains displayed noticeably compromised growth recovery properties compared to that of the WT strain (Fig. 3C). These genes encode an allantoinase (*allB*), a DNA-binding transcription dual regulator (*leuO*), a small, possibly fragmented, protein of unknown function (*ynfP*), and a putative RNA-binding protein (*ybcJ*) (19). As shown in Fig. 3D, although the doubling time of Δ*allB*, Δ*leuO*, Δ*ynfP*, and Δ*ybcJ* mutant strains were similar to that of the WT strain, their lag time to growth recovery was increased by ∼1.5-4 h compared to that of the WT strain. Thus, it seems that the transcription of *allB, leuO, ynfP*, and *ybcJ* during the NSAP are required for optimal growth recovery from long-term N starvation. As the Δ*allB* mutant strain displayed the most pronounced compromised ability to recover growth when compared to the WT strain (Fig. 3C), we decided to focus on the role of *allB* during the NSAP and recovery from long-term N starvation.

**Fig 3.**
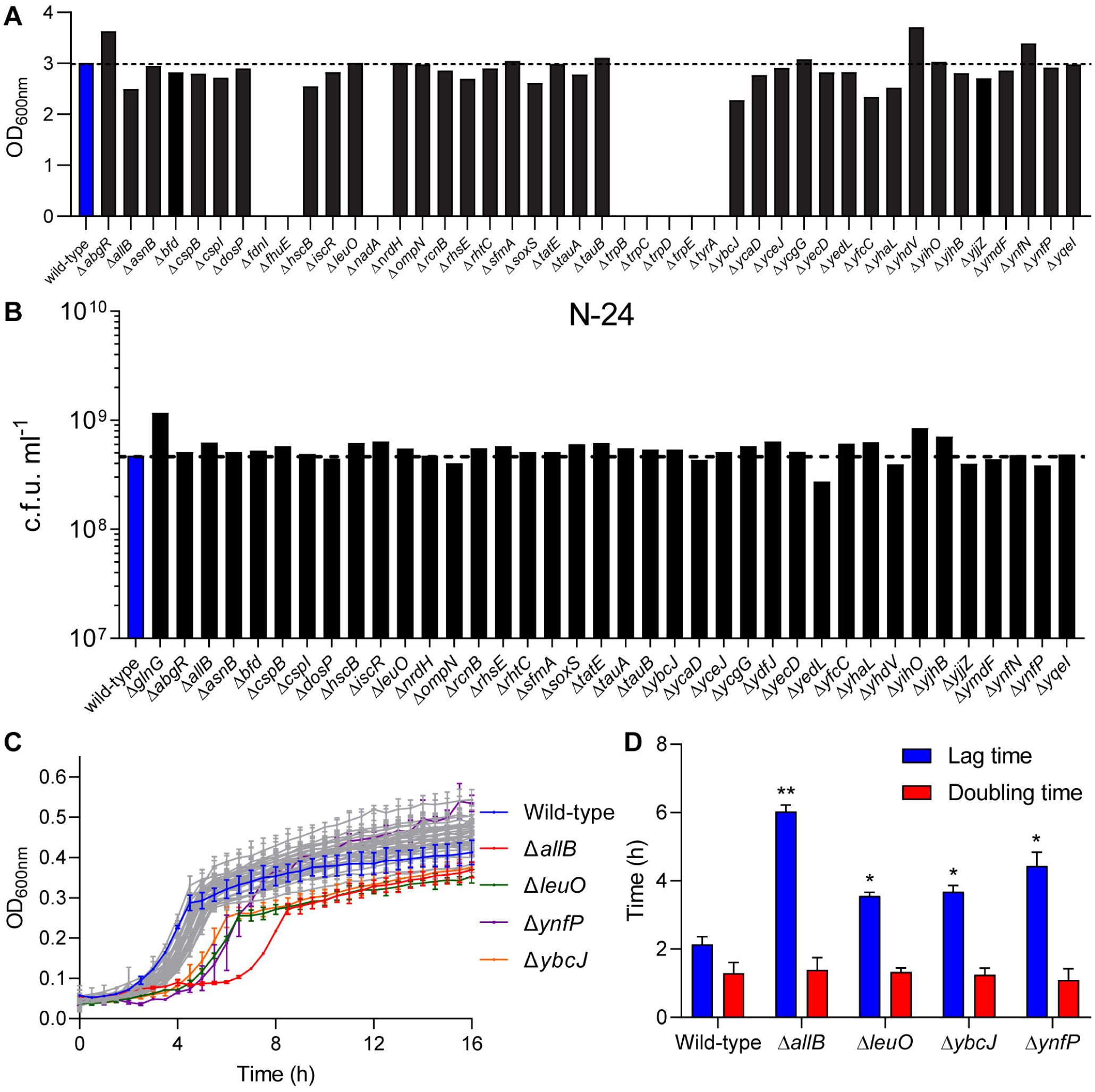
Four upregulated NSAP genes are important for optimal growth recovery from long-term N starvation. (A) Graph showing overnight growth of NSAP gene mutant bacteria as measured by OD_600nm_ compared to WT (blue) bacteria in minimal growth media containing excess (10 mM) N (ammonium chloride). (B) Graph showing viability as measured by c.f.u. counts of wild-type (blue) and NSAP gene mutant bacteria (black) bacteria at N-24. (C) Graph showing recovery growth of 24 h N starved NSAP gene mutant bacteria in N replete fresh media as measured by OD_600nm_ readings. The recovery growth curves of WT (blue), Δ*allB* (red), Δ*leuO* (dark green), Δ*ynfP* (purple), and Δ*ybcJ* (orange) are highlighted. (D) Graph showing lag time (blue) and doubling time (red) of WT, Δ*allB*, Δ*leuO*, Δ*ynfP*, and Δ*ybcJ* bacteria extracted from the growth curves in (C). In (C) and (D), the error bars represent SEM, where *n*=2. Statistical analyses were performed using the Student’s *t*-test (\**P* < 0.05, \*\**P* < 0.01).

### Degradation of allantoin during the NSAP is required for optimal growth recovery from long-term N starvation

To discount the possibility that the difference in lag time during growth recovery between the WT and Δ*allB* bacteria was not due to the presence of a subpopulation of slow growing Δ*allB* bacteria, we measured the lag time distribution of appearance of individual colonies during growth recovery using ScanLag (15) as described in Fig. 1E. As shown in Fig. 4A, we observed the expected lag time in growth recovery for the Δ*allB* bacteria, but no evidence of a slow growing subpopulation was seen. Further, we did not detect a lag in growth recovery of WT and Δ*allB* bacteria sampled from N-when reinoculated into N replete fresh growth medium (Fig. 4B). This result underscores the view that *allB* has a specific role in *E. coli* during the NSAP. To establish that the increased lag time in growth recovery of Δ*allB* bacteria was directly due to the absence of *allB*, we complemented the Δ*allB* bacteria with plasmid-borne *allB*. As shown in Fig. 4C and 4D, the lag time in growth recovery of Δ*allB* bacteria reverted to that of WT bacteria when *allB* was supplied via a plasmid. The product of *allB* is allantoinase: a tetrameric enzyme that catalyses the hydrolytic conversion of (*S*)-(+)-allantoin to allantoate. A previous study (20) reported that substitutions at invariant amino acid positions N94 and S317 (in both cases to D and V), which are located proximal to the active centre, rendered allantoinase inactive (Fig. 4E). As shown in Fig. 4C and 4D, in the context of the complementation experiments, plasmid-borne *allB*-N94V and *allB*-S317V failed to fully decrease the lag time in growth recovery of Δ*allB* bacteria as seen with the WT version of *allB*. In contrast, plasmid-borne *allB*-N94D and to a larger extent *allB*-S317D reverted the lag time in growth recovery of Δ*allB* bacteria as seen with the WT version of *allB*. Although we are unable to explain why specifically substitution by D but not V at N94 or S317 renders allantoinase catalytically inactive under our experimental conditions, it seems that the degradation of allantoin occurs at some point during the NSAP in *E. coli*. This is conceivable because, as shown in Fig. 4F, the conversion of allantoate (the product of allantoinase) to (*S*)-ureidoglycine by allantoate amidohydrolase (AllC) and the subsequent conversion of (*S*)-ureidoglycine to (*S*)-ureidoglycolate by (*S*)-ureidoglycine aminohydrolase (AllE) produces ammonium, which may be needed for maintenance of cellular functions required for optimal growth recovery from long-term N starvation. Intriguingly, compared to *allB*, transcripts corresponding to *allC* and *allE* were only moderately upregulated (∼1.2-1.5 fold) in phases A and B in Fig. 2A and thus did not fall into the category of NSAP genes. However, we observed that 24 h N starved Δ*allC* and Δ*allE* bacteria displayed a similarly increased lag time to growth recovery as the Δ*allB* bacteria (Fig. 4G and 4H) when grown in fresh N replete growth media. This suggests that the activity of the genes of the allantoin degradation pathway have a role in the adaptive response to long-term N starvation in *E. coli*. As allantoin is a product of the purine degradation pathway in N starved *E. coli* (21), we considered whether the requirement for *allB* is a consequence of purine degradation in *E. coli* during the NSAP to sustain cellular functions via the allantoin degradation pathway (as in Fig. 4F). To indirectly determine whether purines are degraded in *E. coli* during the NSAP, we measured the integrity of total RNA as a surrogate for purine degradation in *E. coli* as a function of time in the NSAP by chip-based capillary electrophoresis (see Materials and Methods). As shown in Fig. 4I, strikingly, we observed a gradual degradation of 16S and 23S ribosomal RNA as the NSAP ensued. This observation further underscores the view that the RNA landscape and turnover during the NSAP is plastic and consistent with the view that the transcriptome of *E. coli* undergoes changes during long-term N starvation. Collectively, the results support the view that *allB* contributes to optimal growth recovery from long-term N starvation by the degradation of allantoin during the NSAP in *E. coli*, which likely accumulates as a result of purine degradation.

**Fig. 4.**
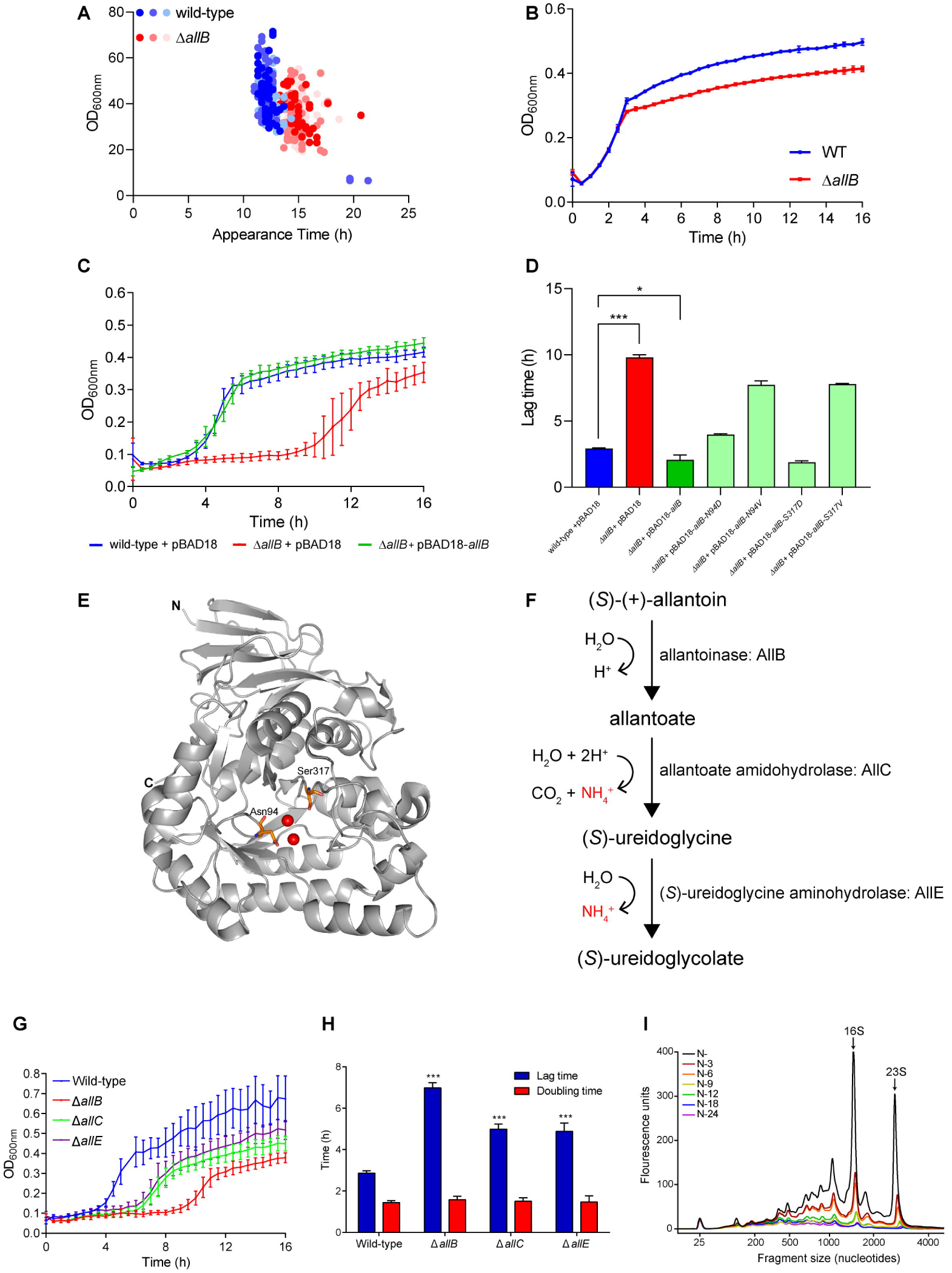
Degradation of allantoin during the NSAP is required for optimal growth recovery from long-term N starvation. (A) ScanLag analysis of colony growth rate and appearance time after plating of 24 h N starved WT (blue) and Δ*allB* (red) bacteria on LB agar. Various shades of the same colour represent distinct biological replicates, where n=3. (B) Graph showing the recovery growth curves of WT (blue) and Δ*allB* (red) bacteria from the onset of N starvation (N-) in fresh N replete media as measured by OD_600nm_ readings of the cultures. (C) Graph showing the recovery growth curves of 24 h N starved WT + pBAD18 (blue), Δ*allB* + pBAD18 (red), and Δ*allB* + pBAD18-*allB* (green) bacteria in fresh N replete media as measured by OD_600nm_ readings of the cultures. (D) Graph showing lag time to growth recovery of WT + pBAD18 (blue), Δ*allB* + pBAD18 (red), Δ*allB* + pBAD18-*allB* (bright green), Δ*allB* + pBAD18-*allB* N94D, Δ*allB* + pBAD18-*allB* N94V, Δ*allB* + pBAD18-*allB* S317D and Δ*allB* + pBAD18-*allB* S317V in fresh N replete media as extracted by OD_600nm_ readings of the cultures. (E) Cartoon representation of the structure of a monomer of AllB in which the positions of the N94 and S317 are indicated in orange relative to the two Fe moieties at the active site of the enzyme (indicated in red). (F) A schematic showing the allantoin degradation pathway indicating the steps where ammonia is produced. (G) As in (C), but the experiment was conducted with Δ*allC* and Δ*allE* bacteria. (H) Graph showing lag time (blue) and doubling time (red) of WT, Δ*allB*, Δ*allC* and Δ*allE* bacteria extracted from the growth curves in (G). (I) Electropherogram of total RNA from bacterial cultures taken at indicated timepoints. Graph shows representative data from at least three biological replicates. 16S and 23S rRNA peaks are indicated. In (B), (C), (D), (G) and (H) error bars represent SEM, where n=3.

## CONCLUSIONS

N is central to all metabolic process in the bacterial cell. As such, bacteria can utilize a wide range of N containing compounds as sole sources of cellular N. Upon N run-out, *E. coli* cells activate the transcription of genes associated with N assimilatory metabolic pathways necessary for utilization of N from the extracellular environment and catabolic biosynthetic pathways for intracellular assimilation of N. We have presented evidence that the growth of a population of N starved *E. coli* does not completely cease upon N run-out and that *E. coli* cells continue to divide over a period of up to 24 h – the NSAP - following entry into N starvation. It seems that the NSAP represents an important adaptive phase in *E. coli* experiencing long-term N starvation to maximise chances of viability and optimise growth recovery should N become replenished. We have also presented evidence that the adaptive transcriptional response to N starvation in *E. coli* occurs in at least two distinct temporally coordinated phases, where the NtrC dependent initial response to N starvation is followed by – to a large extent – a NtrC independent transcriptional response. We have uncovered that an aspect of the adaptive transcriptional response, which occurs in the NSAP subsequent to the initial adaptive transcriptional response in *E. coli* under long-term N starvation, involves the degradation of allantoin. As ammonium is a by-product of degradation of allantoin, we propose that allantoin degradation allows the assimilation of ammonium into glutamine and glutamate to maintain cellular processes during the NSAP to maximise chances of viability and optimise growth recovery when N becomes replenished. Previous studies have shown that allantoin can *only* be used as the sole N source by *E. coli* grown under anaerobic conditions (22, 23), and consistent with these results, we were unable to grow Δ*allB* and WT bacteria under our conditions when ammonium chloride (the sole N source used in our experiments) was replaced by allantoin - in both, the initial growth and recovery growth experiments (data not shown). Therefore, as the end-product of allantoin degradation under aerobic conditions is 3-phosphoglycerate, in addition to the by-product ammonium, we propose that the breakdown of allantoin, which likely occurs as a consequence of purine catabolism during the NSAP in *E. coli* under long-term N starvation, serves to maintain some sort of ‘metabolic threshold’ to allow optimal growth recovery of *E. coli* from long-term N starvation under aerobic conditions when N becomes replenished. In support of this view, the current study and previous work by us (12, 14) and others (13) has also shown that genes associated with the degradation of pyrimidine (e.g. the *rut* operon) become upregulated at the onset of N starvation and remain upregulated throughout the NSAP (14) Thus, collectively, it seems that the degradation of nucleotides is an essential component of the adaptive response to N starvation in *E. coli*. Although future work will focus on deciphering the metabolic and nucleotide landscape to unravel in depth the adaptive responses that define the NSAP in *E. coli*, this work has underscored the importance of studying the transcriptional changes that occur in long-term nutrient starved bacteria to further our understanding of bacterial stress physiology.

## MATERIALS AND METHODS

### Bacterial strains and plasmids, growth conditions and viability measurements

WT and mutant *E. coli* BW25113 strains were used in this study (18). Bacteria were grown as described in Figueira et al (24) in 10 mM NH_4_Cl (for overnight cultures) or 3 mM NH_4_Cl (for N starvation experiments) Gutnick minimal medium, consisting of 33.8 mM KH_2_PO_4_, 77.5 mM K_2_HPO_4_, 5.74 mM K_2_SO_4_, 0.41 mM MgSO_4_, supplemented with Ho-LE trace elements and 0.4 % (w/v) glucose, using NH_4_Cl as the sole N source (16). Where relevant, rifampicin was added at a final concentration of 150 μg/ml at N-1 or N+ as stated, and L-arabinose was added to complementation strains containing the pBAD18 plasmid at 0.2% (w/v) for induction of gene expression in growth cultures. Ampicillin was added at a final concentration of 100 μg/ml to all cultures containing derivatives of the pBAD18 plasmid. Measurement of growth was taken by measuring the optical density (OD_600nm_) of bacterial cultures over time to generate growth curves. Ammonium and glucose concentrations in the media were determined using the Aquaquant ammonium quantification kit and glucose assay kit, respectively, (both by Merck Millipore, UK) according to the manufacturer’s instructions. To determine population viability, the number of viable cells was calculated by conducting serial dilutions of cultures at designated time points as shown in relevant figures, grown overnight on Luria-Bertani (LB) agar plates, and measuring colony forming units (c.f.u.) per ml. The pBAD18-*allB* plasmid was made by traditional restriction enzyme cloning to incorporate the ribosomal binding site of pET33b(+) with the *allB* gene to allow arabinose-inducible expression of the AllB protein. The N94D, N94V, S317D and S317V mutations in *allB* were done by standard site-directed mutagenesis using pBAD18-*allB* as the template.

### ScanLag

Bacteria were grown in 3 mM Gutnick minimal medium to N-24, washed twice in 1 ml PBS, and diluted in PBS. One hundred microlitres of diluted culture was spread on an LB agar plate and incubated at 33 °C for 48 h with pictures of plates taken every 20 minutes. Analysis of colony growth rate and appearance time was modified from Levin-Reisman et al (25) by Dr Miles Priestman and is available at https://github.com/mountainpenguin/NQBMatlab.

### RNA-sequencing (RNA-seq)

The RNA-seq data set was previously generated and analysed by Switzer et al (14). The analysis of differential gene expression during NSAP as well as the statistical analyses were also conducted in the same way as described previously. The sequencing data can be accessed under accession number E-MTAB-6491 in the ArrayExpress database at EMBL-EBI (www.ebi.ack.uk/arrayexpress).

### Total RNA analysis

Cultures were grown as above and sampled at the following time points: at the onset of N starvation (N-) and 3 h (N-3), 6 h (N-6), 9 h (N-9), 12 h (N-12), 18 h (N-18), and 24 h (N-24) under N starvation. At least three biological replicates were taken at each time point. RNA was acquired and stabilized at specified time points using Qiagen RNA Protect reagent (Qiagen, 76526). RNA extraction was carried out using the PureLink RNA mini kit (Invitrogen, 12183025) and DNase Set (Invitrogen, 12185010) as per manufacturer’s protocol. Electropherograms of total RNA integrity were generated using an Agilent Bioanalyser 2100 instrument (Agilent, G2939BA) and RNA 6000 Nano kit (Agilent, 5067-1511) as per the manufacturer’s protocol.

### Statistical analysis and growth calculations

Unless otherwise stated in the text, all growth curves and c.f.u. data show the mean average of three independent experiments and variation calculated by the standard error of the mean (SEM). The student’s t-test or one-way ANOVA was used to determine statistical significance if the probability (*P*) value was < 0.05 (* *P* < 0.05, ** *P* < 0.01, *** *P* < 0.001). Lag and doubling times were calculated by first generating a straight-line graph of the exponential growth section of each growth curve. Lag time was defined as the intersection of the straight line graph with the x axis, while the doubling time was determined during exponential growth (26).

## ACKNOWLEDGMENTS

A Wellcome Trust Investigator award WT100958MA and a Leverhulme Trust project grant (RPG-2017-431) funded this work. J. M. is an MRC funded PhD student.

